# Deep Learning for Alzheimer’s Disease: Mapping Large-scale Histological Tau Protein for Neuroimaging Biomarker Validation

**DOI:** 10.1101/698902

**Authors:** Maryana Alegro, Yuheng Chen, Dulce Ovando, Helmut Heinser, Rana Eser, Daniela Ushizima, Duygu Tosun, Lea T. Grinberg

**Affiliations:** Department of Neurology, University of California San Francisco, San Francisco, CA, USA; Berkeley Institute for Data Science, University of California Berkeley, CA, USA; Bakar Institute for Computational Health Sciences, University of California San Francisco, CA, USA; Julius-Maximilians University Würzburg, Würzburg, Germany; University of Sao Paulo Medical School, Sao Paulo, Brazil; Lawrence Berkeley National Laboratory, Berkeley, CA, USA; Department of Radiology, University of California San Francisco, San Francisco, CA, USA; Veterans Affairs San Francisco, CA, USA

**Keywords:** Machine learning, deep learning, convolutional neural networks, Alzheimer’s disease, histopathology, digital pathology, big data, imaging

## Abstract

Deposits of abnormal tau protein inclusions in the brain are a pathological hallmark of Alzheimer’s disease (AD), and are the best predictor of neuronal loss and clinical decline, but have been limited to postmortem assessment. Imaging-based biomarkers to detect tau deposits *in vivo* could leverage AD diagnosis and monitoring beginning in pre-symptomatic disease stages. Several PET tau tracers are available for research studies, but validation of such tracers against direct detection of tau deposits in brain tissue remains incomplete because of methodological limitations. Confirmation of the biological basis of PET binding requires large-scale voxel-to-voxel correlation has been challenging because of the dimensionality of the whole human brain histology data, deformation caused by tissue processing that precludes registration, and the need to process terabytes of information to cover the whole human brain volume at microscopic resolution. In this study, we created a computational pipeline for segmenting tau inclusions in billion-pixel digital pathology images of whole human brains, aiming at generating quantitative, tridimensional tau density maps that can be used to decipher the distribution of tau inclusions along AD progression and validate PET tau tracers. Our pipeline comprises several pre- and post-processing steps developed to handle the high complexity of these brain digital pathology images. SlideNet, a convolutional neural network designed to process our large datasets to locate and segment tau inclusions, is at the core of the pipeline. Using our novel method, we have successfully processed over 500 slides from two whole human brains, immunostained for two phospho-tau antibodies (AT100 and AT8) spanning several Gigabytes of images. Our artificial neural network estimated strong tau inclusion from image segmentation, which performs with ROC AUC of 0.89 and 0.85 for AT100 and AT8, respectively. Introspection studies further assessed the ability of our trained model to learn tau-related features. Furthermore, our pipeline successfully created 3D tau inclusion density maps that were co-registered to the histology 3D maps.

## Introduction

Compared to other animals, the human brain grew to a considerable size and harbor an astronomical number of neurons and other cells that organized in structures, each one with their particularities and cytoarchitectonic profile. Humans suffer from neurodegenerative conditions that are unique to the species, such as Alzheimer’s and Parkinson’s disease. These neurodegenerative conditions are increasing in prevalence to epidemic numbers, especially with the aging of the population and already cause a high societal and economic burden.

In neurodegenerative diseases (ND), abnormal protein deposits and neuronal loss progressively overtake an expanding landscape of brain areas in stereotypical patterns, providing the neuropathological basis of clinical staging systems. Postmortem examination remains the gold standard for diagnosing, staging and quantifying neurodegenerative disease, a poor substitute for aiding patients during life. However, neuroimaging could potentially facilitate differential diagnosis of NDs, progression monitoring, improve understanding of the underlying pathophysiological processes, and monitor the efficacy of therapeutic agents [1]

Advances in neuroimaging have created new opportunities to detect structural, functional, and molecular brain changes in vivo while preserving the overview landscape. However, no neuroimaging modality offers the resolution obtained by microscopy. Thus the ability to merge different patterns of brain maps obtained at different scales in the same standard spaces would allow merging different information and, as a consequence, improve the analytical power of in vivo neuroimaging. Correlation to histology is considered the gold standard validation method for neuroimaging.

Molecular imaging is particularly relevant to NDs, by allowing spatially-resolved detection of abnormal proteinaceous deposits. [2, 3]. Since abnormal protein deposits start accumulating years before the clinical onset, an ability to reliably image and quantify them *in vivo* would enable earlier case management and vastly improved trials of new potential therapies. Because of this potential, recent years have seen the rapid development of several positron emission tomography (PET) radioligands for *in vivo* labeling of tau and β-amyloid [2, 3]., neuropathological hallmarks of the most common ND, Alzheimer’s disease [4], although as yet only a few β -amyloid tracers are approved for clinical use. Many potential tracers have been years in the experimental stages. Because their accuracy and specificity to detect ND-associated protein deposits remain unclear. While PET-based β-amyloid and tau studies have shown good correlations with postmortem analyses of β -amyloid burden in severe AD stages[5–7], these tracers’ sensitivities to detect scarce protein deposits, remain insufficient [8, 9]. Also, questions about the nature of the off-target binding, the influence of aging, and comorbid pathologies in the signal are still open [10–12] highlighting the importance of developing validation methods that can serve as the basis for improving the interpretation of PET images. However, the key barrier to widely implementing such technologies for diagnosis and staging methods is lack of reliable, validated means to directly assign molecular signals detected by external imaging to the corresponding neural microstructures from which they emanate. High-resolution validation methods allowing for dense and localized (e.g., voxel-to-voxel) comparison of PET signal to histological measures of the target protein and other elements involved in tracer binding (off-target signal), is the stepping stone to expedite the inclusion of more PET tracers in the clinical repertoire. Quantitative 3D mapping of abnormal protein inclusions of the whole human brain has never been achieved because they require adding multiple-step detection methods (immunohistochemistry) in large amounts of brain tissue which generates even more deformation, tissue loss and a method to detect and quantify the abnormal protein in situ that has to resolve differences in shading and background noise. All these also generate even more massive datasets of images. Finally, for assisting validation, these protein maps need to be registered to the neuroimaging datasets with high spatial precision.

Here, we present the solution for quantitative 3D mapping of abnormal protein deposits in the human brain using a semi-automated computational pipeline. Our pipeline leverages our previous solutions for histology to imaging co-registration and introduces a wide range of computer vision techniques together with modern deep learning (DL) algorithms and high-performance computing (HPC) capabilities, being able to process large-scale histological datasets, leveraging a voxel-to-voxel correlation to neuroimaging modalities. It took us three years of full-time work to develop this pipeline. We chose to focus on AD-tau inclusions because most of the current efforts in PET tracer development on neurodegenerative diseases focus on tau

We provide a detailed description of the results and quality control steps of the supporting computational methods of our pipeline. We will illustrate the utility of our pipeline for calculating robust anatomical priors to validate neuroimaging methods applied to two postmortem human brains, harboring moderate and severe Alzheimer’s disease pathology, respectively. The proposed pipeline allowed us to construct quantitative 3D whole human brain maps of tau protein detected by different antibodies at early stages for the first time in the literature. To develop and demonstrate the functionality of the proposed pipeline, we immunostained and digitalized over 500 whole mount human brain slides at high resolution, spanning several Gigabytes of data, and created 3D maps of three different forms of abnormal tau protein in each case and registered each 3D histological map to the MRI volume. Quality control steps showed excellent agreement between DL-based segmentation of tau inclusions against manual segmentation results based on the area under the ROC curve and Dice coefficient metrics. Implementation of these pipeline represents a way forward to validate molecular imaging and advance diagnostics tools for neurodegenerative conditions.

## RESULTS

### Pipeline for creating High-Resolution 3D mapping of immunohistochemical findings in whole human brains

We engineered a comprehensive pipeline (Figure 1) suitable for rendering quantitative 3D mapping of objects of interest (lesions, inclusions) in the whole postmortem human brain that can be deformed for co-registering with other 3D maps of the same brain. Our pipeline relies on three pillars: whole-brain histological processing [13, 14], 3D and 2D registration algorithms [13], and novel deep-learning algorithms to segment and quantify objects of interest in terabytes-large imaging datasets.

**Figure 1:**
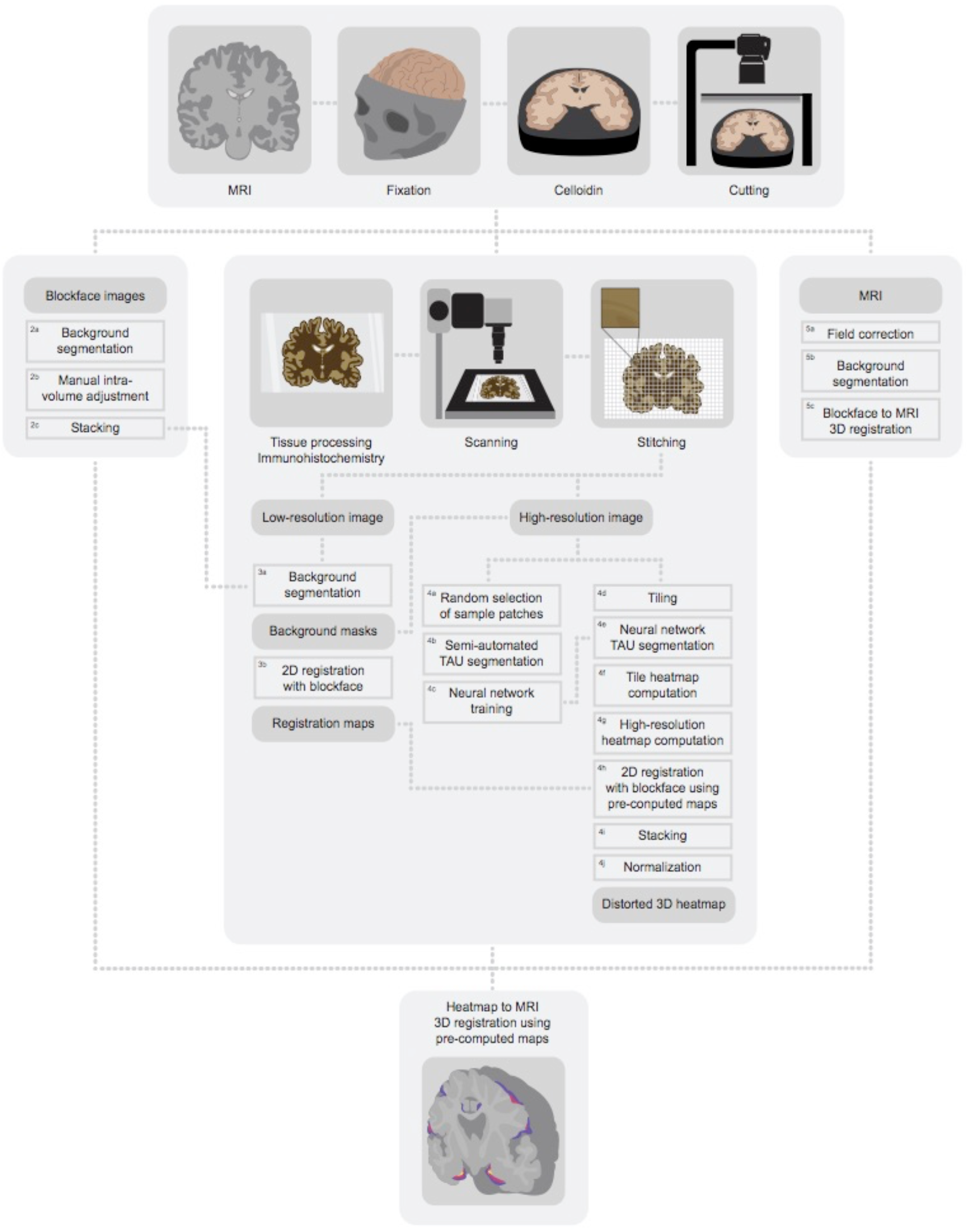
Tau inclusion mapping pipeline. 1. Imaging module; 2. Blockface pre-processing module; 3. Low-resolution histology pre-processing module; 4. High-resolution histology segmentation module; 5. MRI pre-processing module; 6. 3D heatmap to MRI registration.

We previously had developed and published the protocols for the two first pillars. Here, we will focus on the pipeline development and quality control step for enabling scaling whole -brain section immunostaining and developing the deep learning algorithms for segmentation. Because of the scale of the human brain, <u>we had to create our imaging equipment and software to enable analysis of such large histological</u> sections. To demonstrate the clinical usefulness of our pipeline, we chose to test it for quantifying and mapping various abnormal tau proteinaceous inclusions seen in Alzheimer’s disease. We processed two whole human brains (one with moderate and another with severe AD neuropathology - case #1: Braak stage 4 [15] and case #2: Braak stage 6) in 1608 coronal whole brain sections of which 524 underwent immunohistochemistry. Then, we used our computational pipeline to scan the histological sections at microscopic resolution, quantify tau inclusions in each voxel, map the inclusions in 3D and register those maps to the corresponding MRI volume.

We broke down our pipeline in a series of histological and computation modules (Figure 1) that can be executed independently to allow for scalability. Briefly, the first module includes acquiring structural MRI to create an **MRI dataset**(<u>*Neuroimaging Acquisition Module).*</u> We acquired postmortem MRI sequences in cranio, right before procurement [13]. Next, upon procurement, the brain was processed and embedded as a whole in celloidin and coronally cut into 160-micrometer thickness serial sections [13, 16] (<u>*Histological Processing Module*</u>). The section thickness was selected to balance the risk of tissue tear and the efficiency of antibody penetration. Following the brain slicing process, the pipeline created two additional datasets: (a) **serial histological sections**(about 800 per case, coronal axis) followed by the <u>*Immunostaining module*</u> and, (b) **blockface image dataset** comprising digital images of each histological section generated by photographing the celloidin block before each microtome stroke with a camera mounted on a photo stand. The **blockface image dataset** proceeds to the <u>*Blockface Processing Module,*</u> which generates a *blockface volume*, which is an intermediate space for registering both the 2D tau maps and the MRI volume for enabling neuroimaging to histology voxel-to-voxel comparisons [13]. We provide details of the <u>*Neuroimaging Acquisition, Histological processing, and Blockface processing modules*</u> in Methods and in our previous publications [13, 14, 17’19]. Hereon, we will focus on the Modules of this pipeline developed specifically for this study.

It is often unclear what is the biological basis of the signal or which form(s) of a protein a PET-tracer is binding in vivo. Therefore, probing different histological targets within the same image voxel is important for validating neuroimaging results.

Alzheimer’s disease courses with the accumulation of different types of abnormal tau protein. Using immunohistochemistry in the <u>*Immunostaining module*</u> (Figure 2), we generated histological section sets to detect three different abnormal tau forms per case (6 maps in total). We labeled phospho-tau Ser 202 with monoclonal antibody AT8, phospho-tau Ser 404 with monoclonal antibody AT100, and conformationally changed tau, recognized by monoclonal antibody MC1. Clinical neuroimaging offers a spatial resolution spanning from 1 to 5 mm. Conversely, the spatial resolution of histological sections is much higher, creating an opportunity to generate multiple histological maps within one image-voxel. Using our protocol, a series of 4 histological sections encompass a 1mm thickness after registration to MRI. Thus, we divided the histological sections into approximately 200 sets of 4 sections to accommodate the different tau stainings at an interval in which each image voxel would have a corresponding histology labeling for each antibody and still save extra sections for additional or repeat stainings. Each whole-brain coronal section selected for immunostaining was immunolabeled with one antibody only, to avoid mixing signals, and mounted on 6” x4” inch glass slides. We maximized the number of sections per staining batch to avoid bias due to inherent batch variations. A single technician could immunostain two batches of 24 sections per week. Thus, it takes about four weeks to immunostain sections corresponding to a 1 mm interval (200 coronal sections if spanning the whole brain). Details of the immunohistochemical method, including adaptations for enabling staining whole coronal sections, using free-floating protocols are in *Methods*.

**Figure 2:**
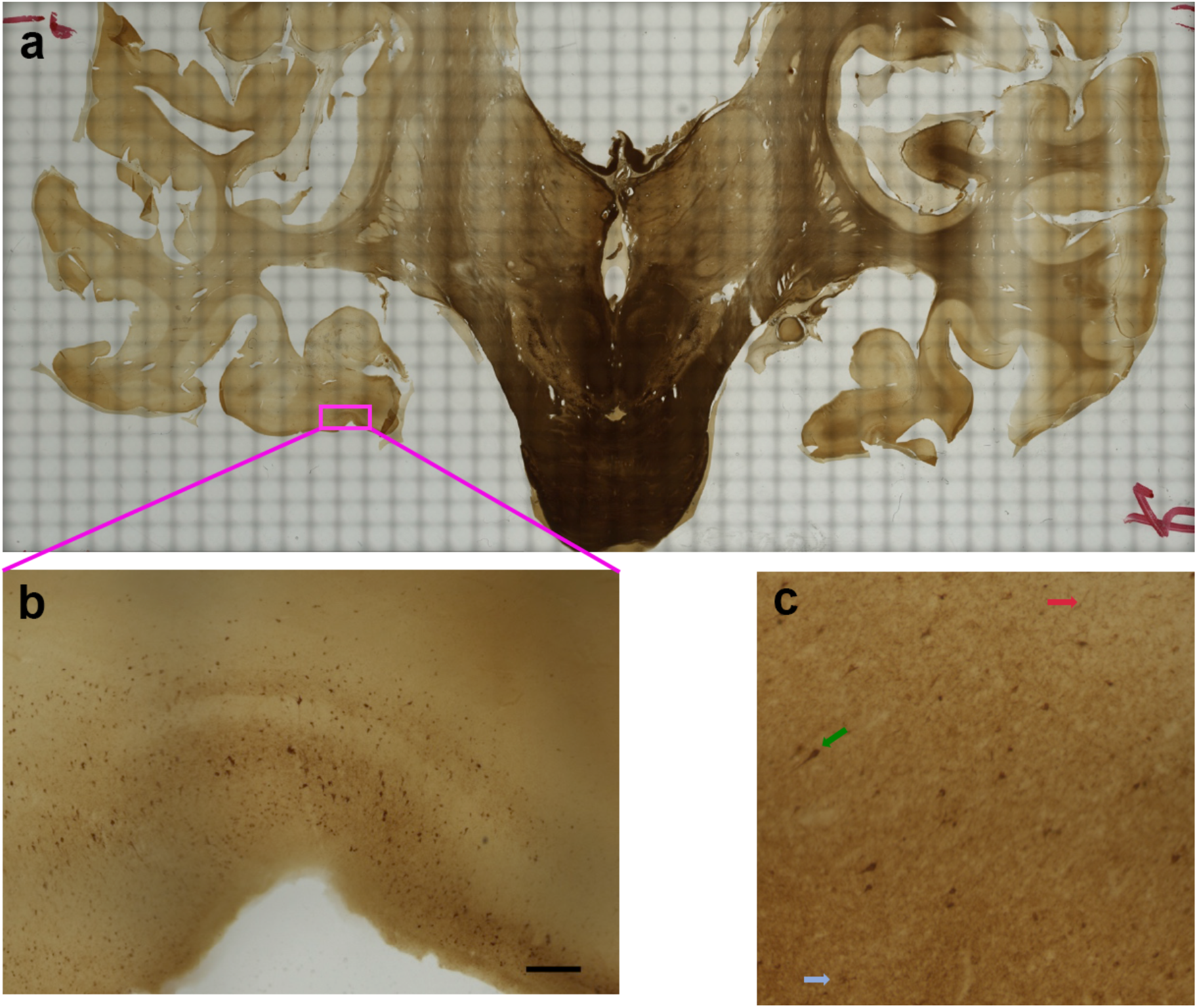
a) example of a stitched image; b) 100% zoom of a hippocampus area (scalebar is 0.3mm); c) example of tau inclusions our system is capable of detecting. Green arrow shows an NFT, red arrow shows a thread and blue arrow show as plaque.

We designed a custom-built whole-slide scanner (Figure 3) because no commercially-available option was available to accommodate large tissue sections and required microscopic resolution we aimed to achieve. A total of 524 immunostained sections were scanned for quantifying tau inclusions (<u>*scanning and stitching module).*</u> The scanner hardware features a high-precision industrial XY stage, a color CCD camera, and a 5.5X machine vision objective mounted directly on the camera. As the objective field of view has 3.28 × 2.6 mm, scanning a whole-brain section required stitching hundreds of images tiles. We developed an in-house software using Macro Manager 2.0 to control the scanner. The software has a user-friendly interface that allows the user to select the region-of-interest (ROI) for automatic computing the coordinates of the image tiles necessary to cover the selected ROI and synchronize the XY stage movements with image capture. The software also allows for adjusting white balance and magnification parameters of the lens. Each section generated about 1500 image tiles (20GBs of data) at a resolution of 1.22μm/pixel after 90 min of scanning. We used TeraStitcher [20], which is capable of working with several Gigabytes of data while maintaining a small memory footprint stitch the tiles. Since it took approximately two days to stitch each section, and we used an HPC system to perform stitching in an embarrassingly parallel way. The stitching process generated two new imaging datasets. The **high-resolution digital section dataset** with images of approximately 80000×50000 pixels per tissue section (another 20GBs of data per slide) were registered to the blockface volume (Figure 2) and segmented for tau inclusions to generate first 2D and then 3D spatial heatmaps. As it is unfeasible to directly manipulate large datasets in regular workstations, we also created a (b) **low-resolution digital section dataset** in which the original digital sections were downsampled to a 10% resolution to facilitate all computing steps that did not require microscopic resolution, including visual inspection of the final stitched images, 2D registration of the histological imaging to the blockface image (<u>*2D histology registration step)*</u> and masking background and any brain area of interest (Fig1-1b).

**Figure 3:**
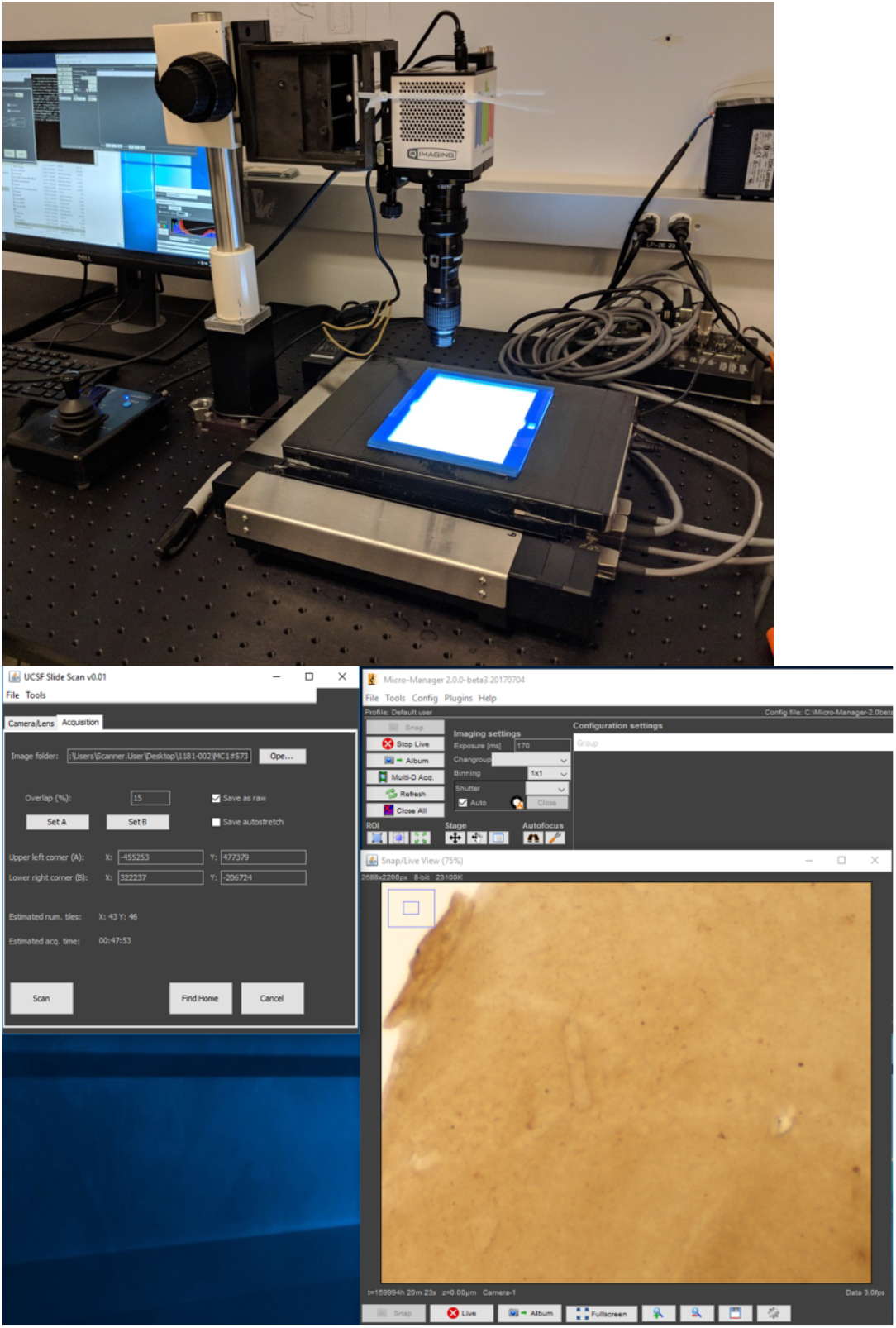
Left: our customized whole slide scanner. Right: User interface of the controller software developed using Micro Manager 2.0

### 2D Inclusion Segmentation And Mapping Module

For simplicity, we broke down this module in sequential steps necessary to prepare the images for training and application of the convolutional neural network (CNN) based DL models. The validation steps will be described in a separate section. CNN-based segmentation is computationally intense. Thus, we first identified imaging regions of interest to decrease computing time. Specifically, we masked out pixels of background and white matter tissue to focus in the imaging regions with tau deposition. Masks were created using a <u>low-resolution digital section dataset</u> and subsequently upsampled and applied to the high-resolution histological images.

#### SlideNet, a convolutional neural network (CNN) for semantic segmentation (Figure 4)

##### Architecture and rationale

Based on the literature, the density of tau inclusions in each pixel could be estimated by applying thresholding algorithms, assuming the immunostaining would develop in brown color in the target tissue (DAB), and the tissue background would be almost transparent. However, despite all optimizing steps during immunohistochemistry to achieve a high signal to noise ratio, the failure rate for this thresholding strategy high for various reasons. First, a conventional 8-um thick histological section is rather transparent, but a section 16x thicker is opaque, especially for the white matter tissue. Second, factors hard to control in an experimental setup such as minimal thickness variations and hydration level influence the shade of the staining in large sections. Finally, because the drying process of such thick sections takes weeks, some degree of microscopic dust may deposit over the tissue. To address these challenges, we designed **SlideNet**, a convolutional neural network (CNN) for segmenting inclusions of interest in a setting of lower signal to noise ratio. **SlideNet**, part of 2*<u>D inclusion segmentation and mapping Module</u>* (Figure 1), uses a UNet based architecture (Figure 4), trained to compute a confidence map of the 2D distribution of an object of interest (i.e, tau inclusion) instead of a single class confidence value. Furthermore, we extended SlideNet architecture to leverage color information and to optimally work with histological imaging resolution (1.22 um/pixel). SlideNet is the core innovation of our pipeline, with higher segmentation accuracy and robustness to histological artifacts compared to thresholding algorithms.

**Figure 4:**
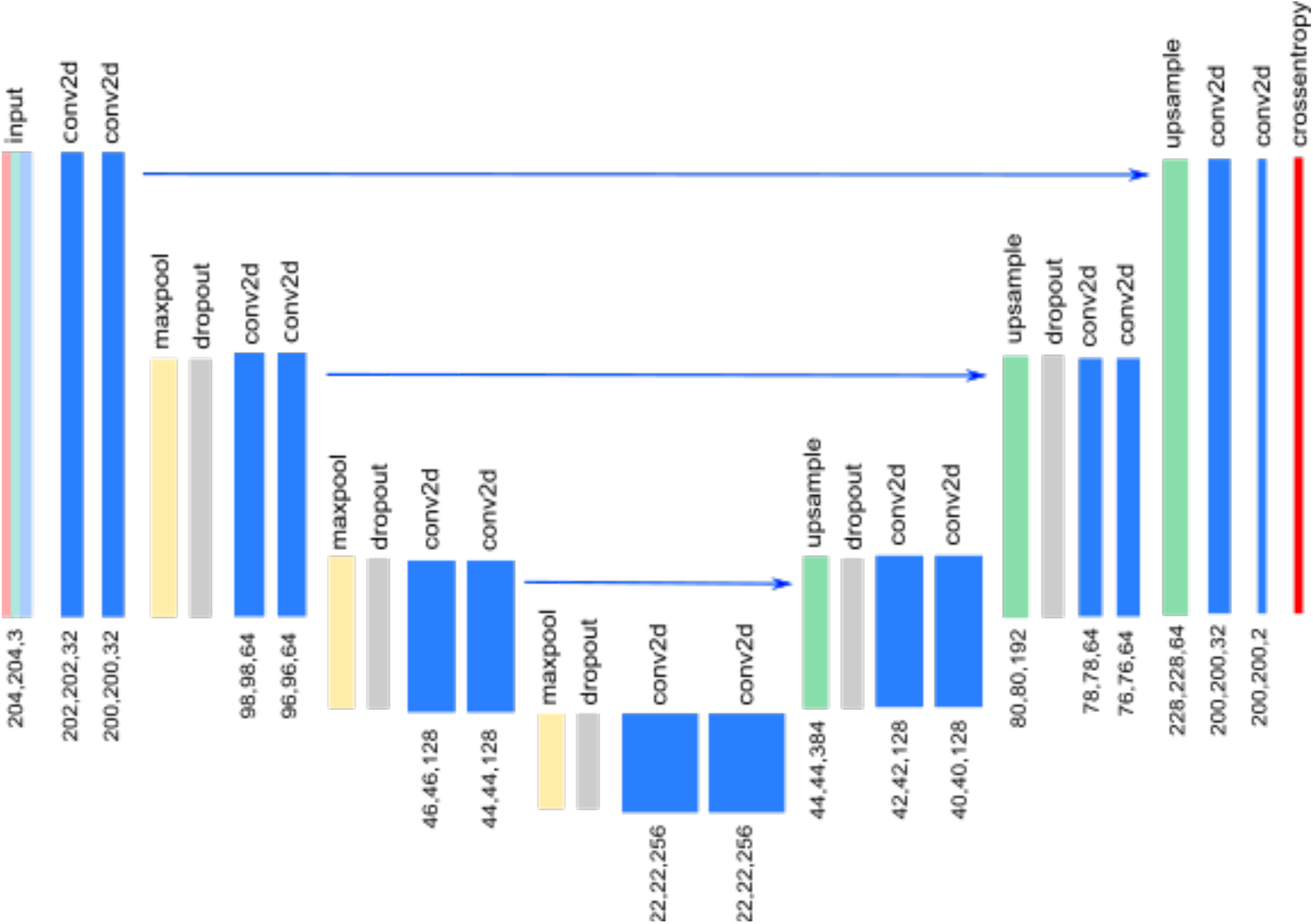
SlideNet architecture. The arrows indicate skip connections that are followed by cropped when the right-hand side layer is smaller than the left side layer

SlideNet has an input layer measuring 204×204×3 pixels that correspond to 1mm^2^ of tissue in our scanner resolution to accommodate RGB images. SlideNet features a validation hold-out scheme, where training and testing sets were used for model training, and a validation set was used for independent network performance statistics.

##### Network training

For each brain, we randomly sampled 100 imaging patches (1024 × 1024 pixels) per antibody (AT8, AT100, and MC1), yielding 600 patches (200 per antibody) from the masked high-resolution histological images. Next, the patches underwent semi-automatic labeling of tau and background by trained observers, followed by visual quality control and manual editing by other observers (LTG) with extensive experience in histological studies. Labeling took approximately 2 hours per patch (total 1200 h or 150 days of specialized work). We used 80% of the patches for training, 10% for testing, and 10% for validation. SlideNET training took up to 2 days per antibody using a workstation with 2 GPUs. The networks were trained until we observed a plateau in the accuracy and loss function graphs. The network output probability maps for each patch, which were thresholded at the cut off of 0.7 to create binary tau inclusion masks. The threshold cut-off was set heuristically, aiming to achieve high precision.

##### Validation of the network training

Aiming to validate the SlideNet model, we computed the receiver operating characteristic (ROC) and precision-recall curves over the testing and validation sets for each antibody. For ROC, the AT8 model achieved an area under the curve (AUC) of 0.85 for both testing and validation datasets, while the AT100 model achieved an AUC of 0.87 on testing and 0.89 on the validation dataset. MC1 model achieved the best ROC results with an AUC of 0.91 on the testing and 0.88 on the validation dataset (Figure 5). From thresholded masks, we calculated precision and recall values. We obtained precision values of 0.87, 0.87, 0.90 respectively for AT8, AT100, and MC1 in the testing datasets and 0.92, 0.92, 0.91 respectively for AT8, AT100, and MC1 in the validation datasets. As for the recall, the thresholded masks obtained 0.16, 0.19, 0.12 respectively for AT8, AT100, and MC1 in the testing datasets and 0.18, 0.13, and 0.08 respectively for AT8, AT100, and MC1 in the validation datasets.

**Figure 5:**
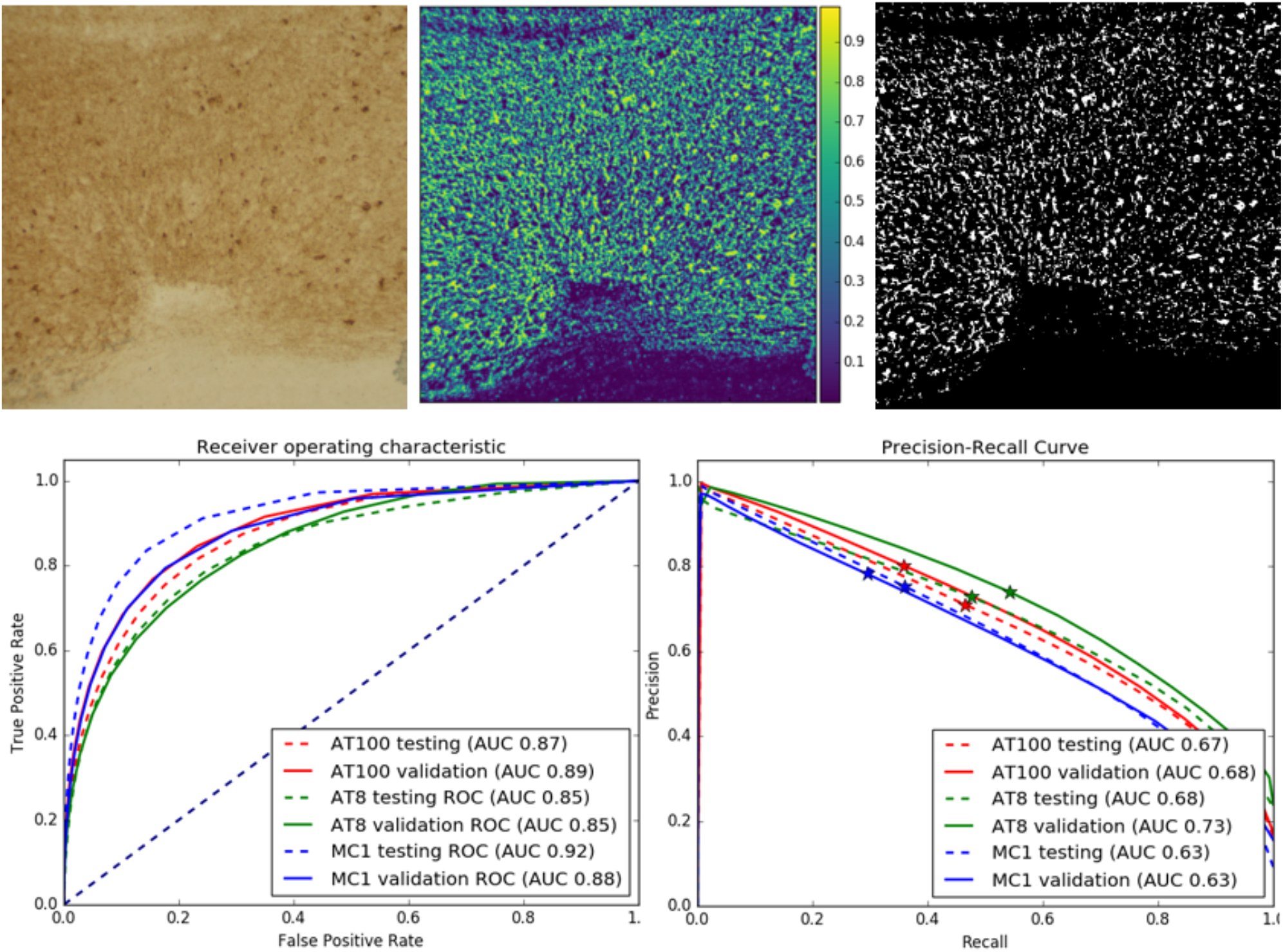
Top row: AT8-stained image patch (left), its respective probability map (middle) and binary segmentation after thresholding at 0.5 (right). Bottom row: graphs show ROC (left) and precision-recall curves (right) for models trained with all full datasets from all stains. Stars on the precision-recall graphs show where the 0.5 threshold is localized for each stain.

In terms of precision-recall, AT8 model achieved an AUC of 0.68 for the testing and 0.72 for validation; AT100 had a 0.66 for testing and 0.68 for validation, and MC1 had a 0.63 for testing and 0.63 for validation.

##### Tau Segmentation of the full resolution dataset

To facilitate computation, we divided the high-resolution images into tiles. We then extracted the tiles corresponding to unmasked areas (i.e., gray matter tissue) and further divided them into 2550 patches of 204 × 204 pixels (i.e., CNN training input image size) with 90 pixels of overlap between sections. It took about four days for SlideNet to segment the largest dataset (AT8, case 2,160 sections). The network output probability maps were thresholded to create binary masks. The same cut-off values for thresholding were applied to all datasets. Figure 5 shows an example of histology tile (left), the probability map created by the CNN (middle), and the thresholded binary mask (right).

##### Tau density maps (heatmaps)

Heatmaps were computed for each tile binary mask (Figure 1f) by calculating the average surface area occupied by tau inside 1μm^2^ of tissue. Next, heatmap tiles were stitched together to form a whole section heatmap (Figure 6).

**Figure 6:**
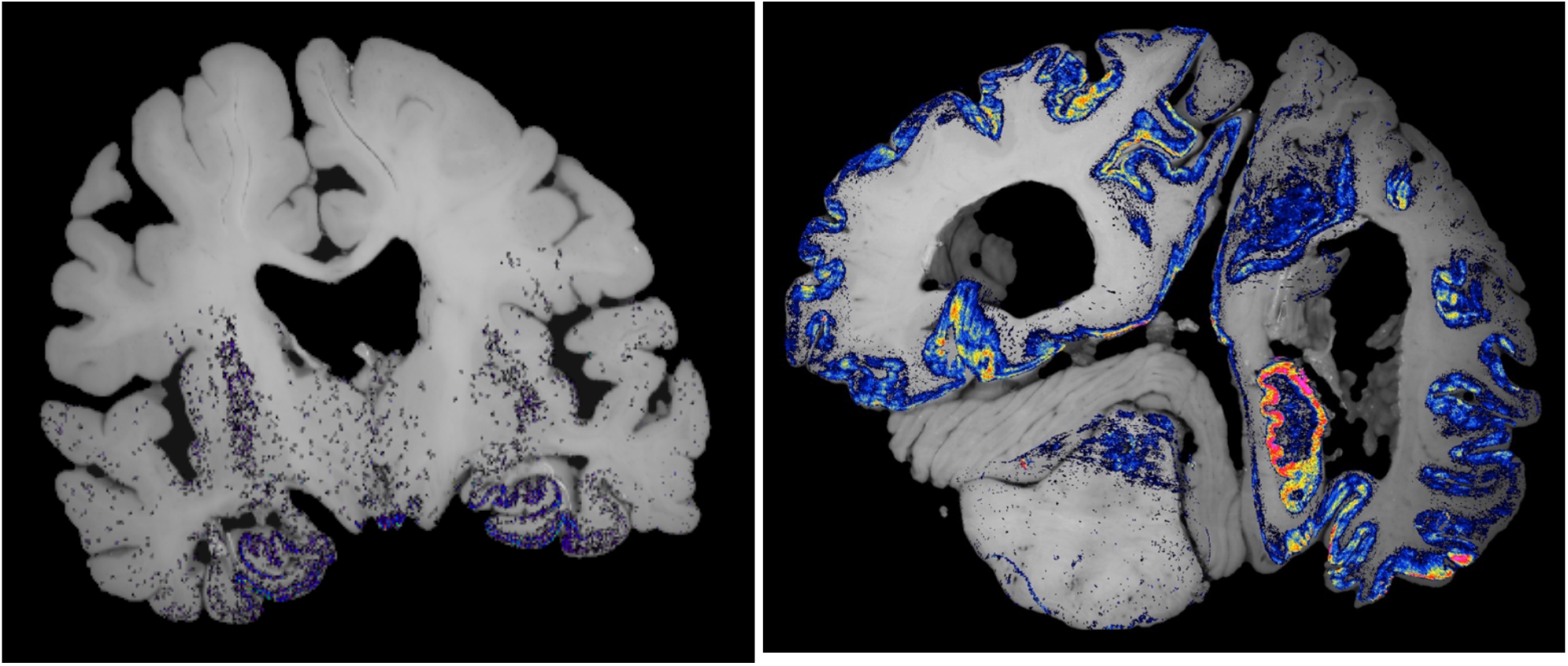
Examples of a heatmap from the early (left) and late (right) stage specimens. Here, warmer colors indicate a higher density of tau tangles.

### 2D histology registration and 3D reconstruction Module

During immunostaining, alcohols and solvents cause non-linear deformation to histological sections. Thus we registered each low-resolution histological digital image to the corresponding blockface image. First, we applied the previously created background masks to the <u>low-resolution digital section dataset</u>. Then, we used a combination of open-source registration algorithms to align these masked images to their corresponding pre-processed blockface images to create warping maps for each section. The 2D registration results were visually inspected, and imperfections were corrected manually—most of the imperfections located around the limbic structures. Finally, we used the resulting warping maps to register each whole section heatmap to their corresponding blockface image, thus allowing for 3D reconstruction of the tau inclusion density maps in the blockface image space. The final 3D tau inclusion maps were stored in standard nifti medical imaging file format-. We rendered 3D map visualizations using Amira. Figure 7 shows examples of early and late-stage heatmaps overlaid on their corresponding blockface images. Here, the hotter colors indicate a higher density of tau inclusions.

**Figure 7:**
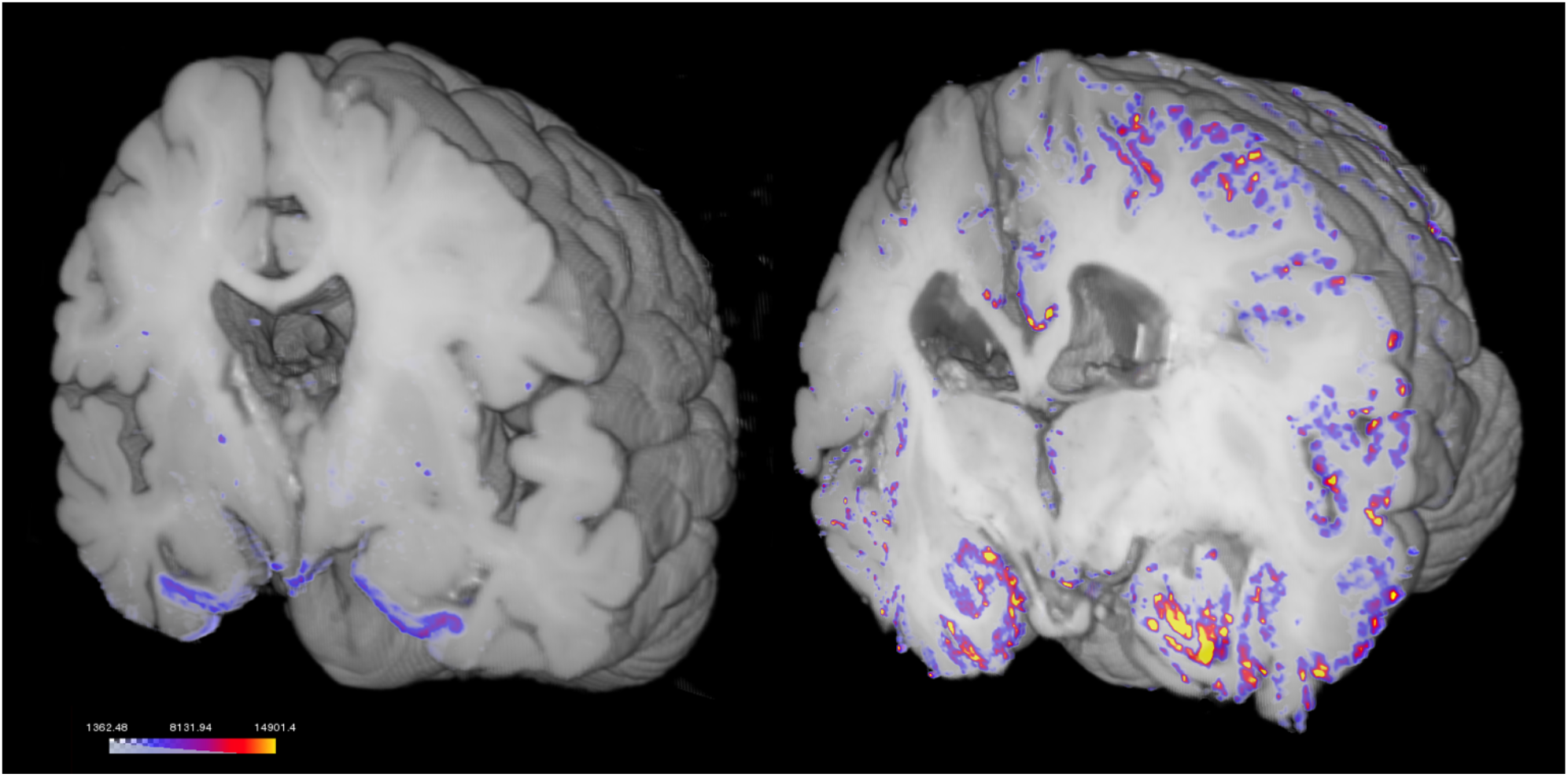
Volumetric reconstruction of the early (left) and late-stage (right) AT100 tau heatmaps around the hippocampus area, overlaid on their respective blockface volumes

#### Histology to MRI 3D registration

The final product of our pipeline was six 3D tau inclusion datasets, showing the distribution of tau stained for AT8, AT100, and MCI, on two different whole human brain samples. Figure 8 shows example volumetric renderings of tau inclusion maps overlaid on their corresponding MRI images on the blockface volume space. Briefly, structural T1-weighted MRIs were warped into the 3D image space of the corresponding blockface reconstructions using diffeomorphic spatial normalization in ANTs [21, 22]. The diffeomorphic transform matrices were then applied to all the tau inclusion heatmaps, for mapping into the structural MR imaging space and resolution.

**Figure 8:**
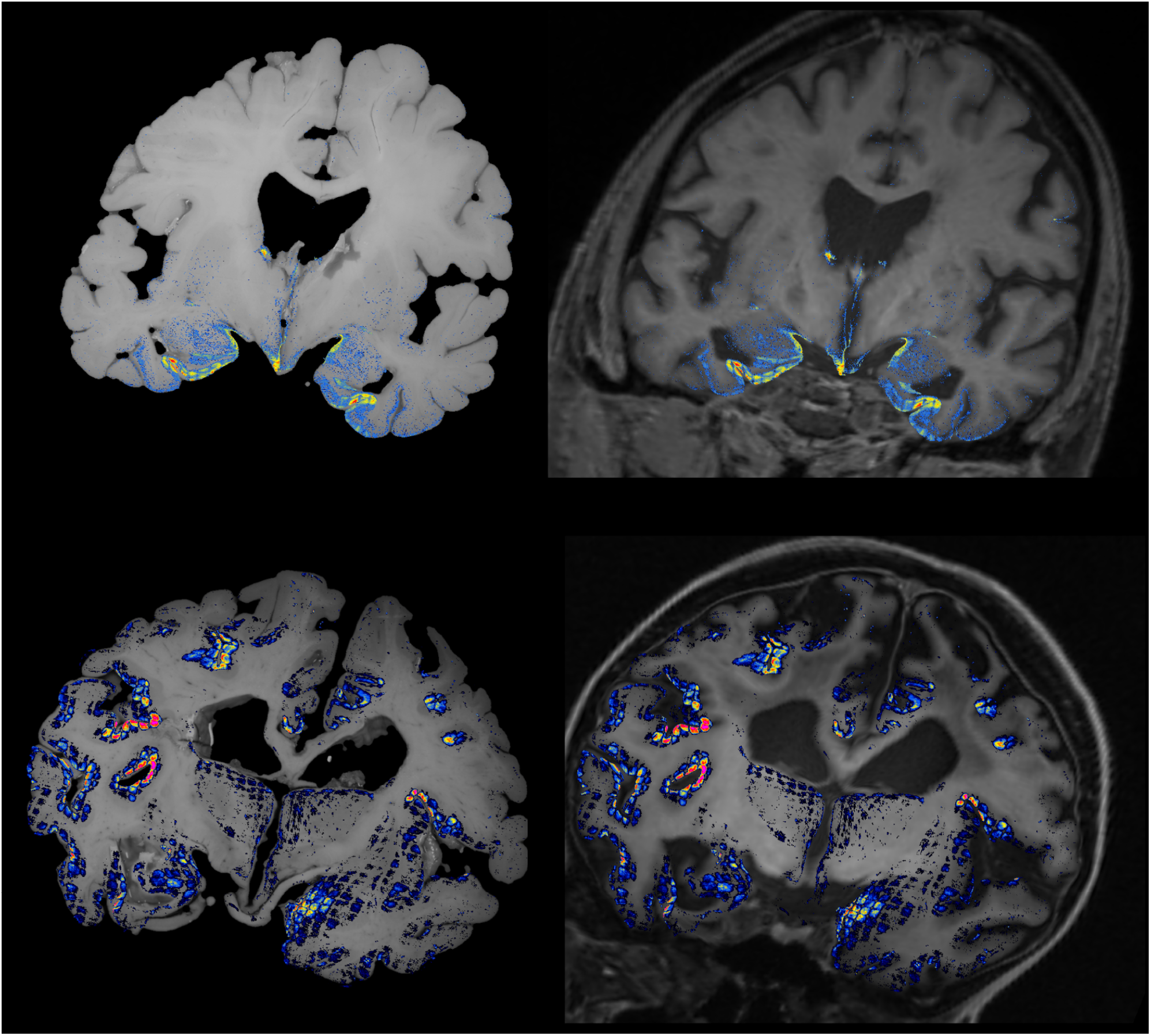
Top row: case #1 heatmap overlaid on blockface (left) and MRI image (right); Bottom row: case #2 heatmap overlaid on blockface (left) and MRI image (right).

### Transfer learning and additional validation experiments

Considering it is unfeasible to dedicate the same amount of effort we used to generate training data for each antibody and case for every new case, we run additional experiments to test how the network behaves with different training datasets size and composition. <u>For these experiments, we used the following training datasets three</u> full datasets as described above (each containing all 200 semi-automatically labeled patches per antibody); 3 case #1 (restricted) datasets (each containing all 100 semi-automatically labeled patches per antibody from case #1); 3 case #2 (restricted) datasets (each containing all 100 semi-automatically labeled patches per antibody from case #2), totalizing nine trained models.

Next, we wanted to test how a network trained with labeled patches from a given antibody from one case, performs in another case. For instance, the network trained using AT8 patches from case #1 segmented AT8 signal on case #2 with an AUC of 0.7855 in testing dataset segmentation and 0.7847 in the validation set. Inversely, when the network trained on AT8 patches from case #2 was applied to segment AT8 signal on case #1, we obtained an AUC of 0.8642 in the testing set and 0.7544 in the validation set (Figure 9).

**Figure 9:**
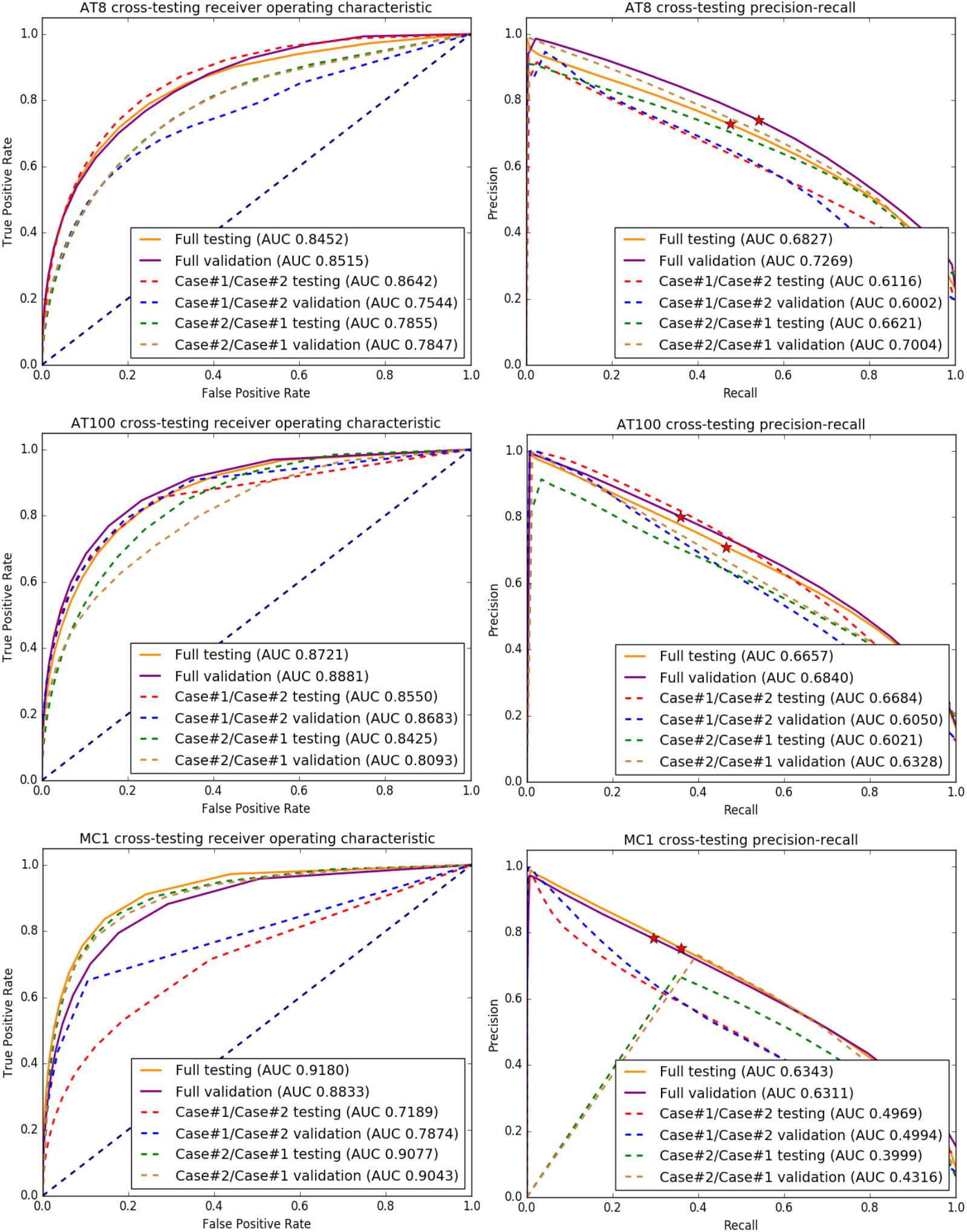
Comparison of ROC and precision-recall curves for models trained with complete datasets (all images from each marker) versus models trained with a dataset comprised of images only from one marker and one brain. <u>Top row</u>: ROC and precision-recall curves for the model trained with the complete AT8 dataset where the solid orange and solid purple lines show the results for testing and validation datasets. Dashed red and blue lines show testing and validation results for the model trained with AT8 images from case #2 alone and dashed green and brown lines show testing and validation results for the model trained with AT8 images from case #1 alone. <u>Middle row</u>: same as top row but using AT100 images. <u>Bottom row</u>: same as top row but using MC1 images.

Finally, we used labeled patches of a given antibody to train a network and then applied the network to a validation dataset of another antibody (cross-testing). When a network trained on patches with AT100 antibody for training was applied to segment AT8 datasets, we achieved an AUC of 0.8459 in AT8 the testing dataset segmentation and 0.8219 in the AT8 validation dataset segmentation. Inversely, a network trained on AT8 patches when applied to segment AT100 datasets resulted in an AUC of 0.8677 in the AT100 testing dataset and 0.8501 in the AT100 validation dataset (Figure 10).

**Figure 10:**
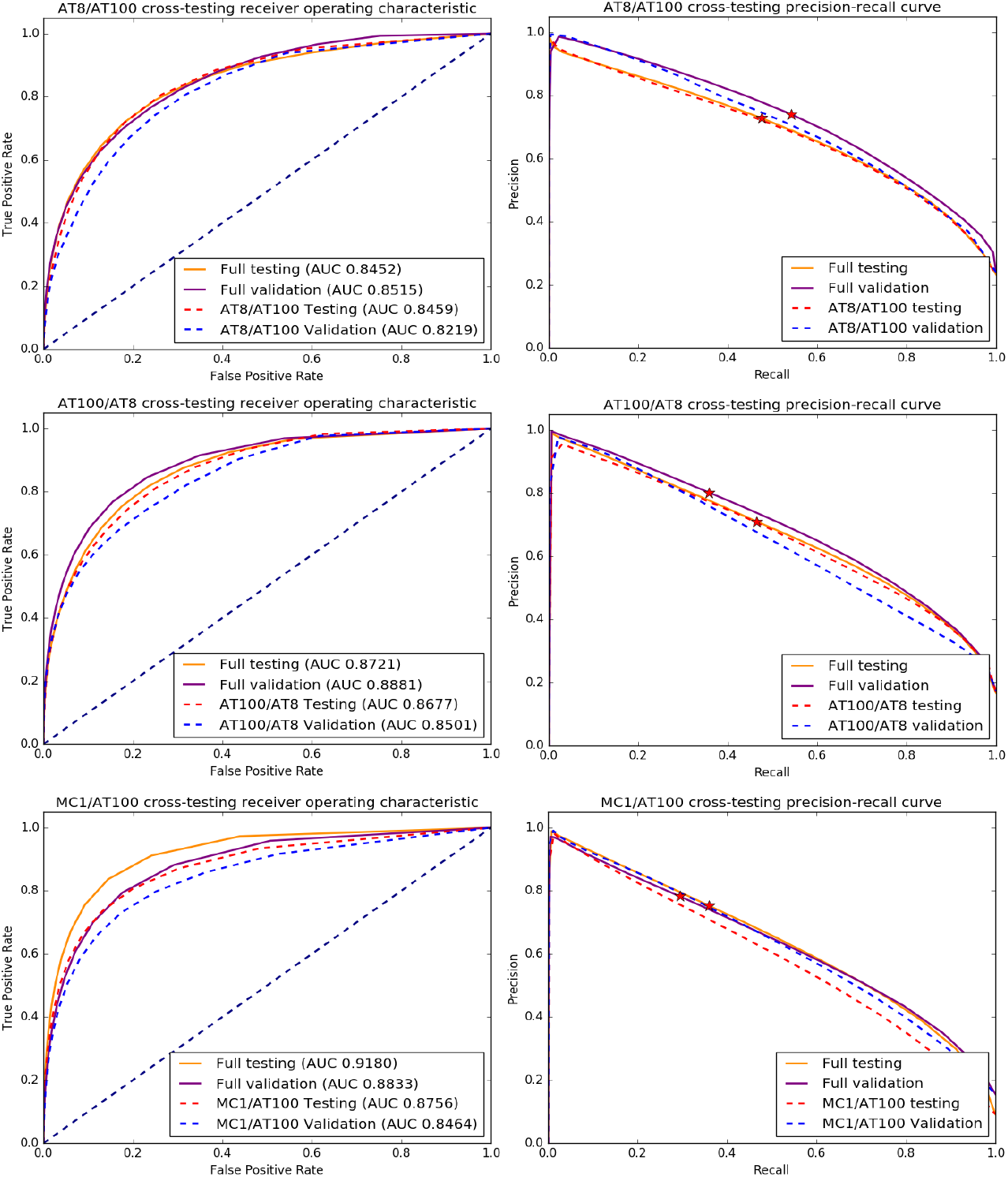
Comparison of ROC and precision-recall curves for cross-trained models, where we performed training using images from one marker and tested the model on a dataset of a different marker. <u>Top row</u>: dashed red and blue lines show results test, and validation results for a model trained using AT100 data and used for segmenting AT8 images. For comparison, the orange and purple lines show the test and validation results for the model trained with AT8 data. <u>Middle row</u>: same and top row, using a model trained on AT8 images to segment AT100 images. <u>Bottom row</u>: same as top row, using a model trained on AT100 images to segment MC1 images.

Again, we see that the difference between models is mostly negligible, with the curves for the models trained with case 1 AT8-only and AT100-only images being slightly inferior to the model trained with the complete dataset shows the result of the cross-training experiment. While the ROC AUC values do not show a huge difference between models, the precision-recall curves show a decrease in model accuracy. These results indicate that mixing images from different Braak stages – as in the full model training where images from cases with different Braak are combined, not only does not interfere with training but is actually beneficial, suggesting that it is possible to use transfer learning [23] thus allowing us to perform the networking training by adding new images and improving on top of a previously trained full model.

#### Network introspection experiments show the neural network correctly learns tau inclusions

Standard precision-recall and ROC curve evaluation are unable to show whether the network is effective learning image features of objects of interest (tau inclusions) or simply overfitting patterns. Thus, we used neural network inspection techniques to evaluate the network ability to learn. We first used gradient guided class activation maps (Grad-CAM) [24] on randomly selected images to evaluate what features were most important to drive the network to a particular decision. Then, we added perturbations for checking whether the network was segmenting tau inclusions solely based on spatial localization, or it was using relevant features.

Standard Grad-CAM works by computing the gradient between a user-defined class neuron in the last network layer and an intermediary target layer and then multiplying its mean value with the target class activations. This technique was designed to work with networks that have a fully connected or a global pooling layer [25], which are responsible for mapping the information spread across the convolutional layers to a single neuron at the last layer. As SlideNet rather outputs 2D probability maps of inclusions and lacks the property of fragmented mapping information to a single neuron, we adapted Grad-CAM by selecting multiple output neurons inside a tau inclusion, on randomly selected test images, i.e., neurons that should be activated for the class “tau,” with the help of a binary mask, and computing the mean of the Grad-CAM maps generated for each selected neuron. As seen in Fig 11, a visual inspection of the output CAM maps shows a good agreement between the features that drove network decision and tau signal, and the network correctly responded to perturbations (Fig 11). We repeated the same experiment to interrogate if the network could correctly discriminate the background. We used a mask to select background neurons, as shown in Figure 11d. The resulting Grad-CAM (Figure 11e), shows good localization of the background as well. Finally, we interrogated if the network was learning to detect tau based on pixel spatial localization only or was using other relevant information. We created a perturbed image by partially covering the Tau tangle with a patch of background and repeated the background Grad-CAM experiment (fig 11f/g). The network correctly recognized the patch as background, suggesting that other relevant features inform the network.

**Figure 11:**
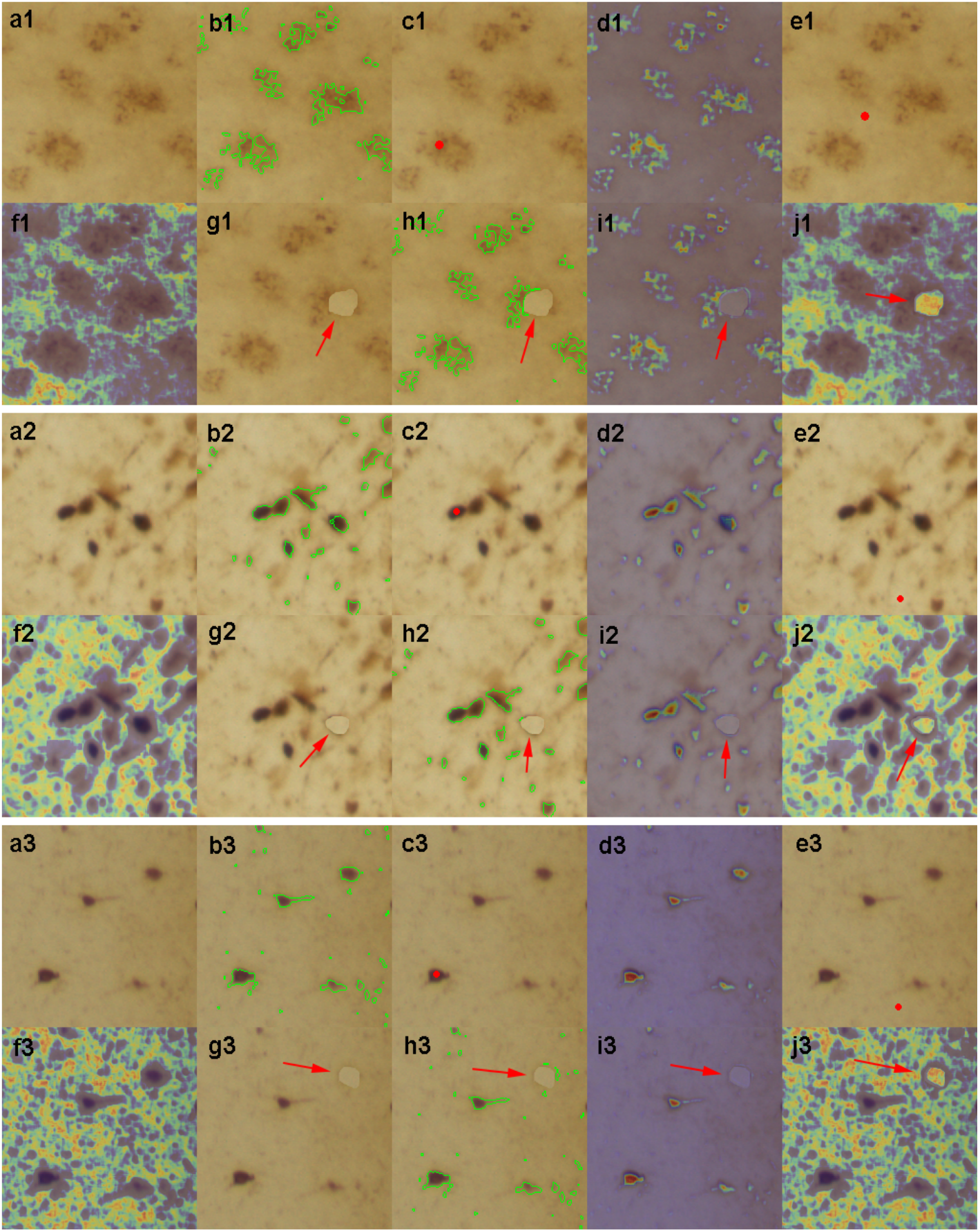
Network interpretability using Grad-CAM and perturbation techniques. a. Example of original tau tangle with some threads. b. Mask used for selecting pixels from (a). c. Grad-CAM result using pixels from mask in (b) as reference. d. mask used to locate background pixels. e. Grad-CAM results using background pixels as reference. f. Perturbed image, where a background image patch (indicated by the red arrow) was artificially placed to partially cover the tau inclusion. g. mask used for locating background pixel on the perturbed image. h. Grad-CAM results using the perturbed image. The red arrow indicates the perturbed region that was correctly considered background. The result can be seen in Figure 8, where Figure 8a shows the original test image, 8b the binary mask, and 8c the estimated Grad-CAM, with brighter values indicating stronger influence in the network decision.

### Registration validation

We quantitatively evaluated registration quality by computing the Dice coefficient (DC) [26] between stacks of masks manually labeled on MRI and histological slices. The Dice coefficient measures how well co-localized two sets of images are, being a popular image registration metric.

#### Histology to blockface 2d registration evaluation

we manually labeled the superior ventricles on histological images for all tau markers, excluding images where the ventricles were not completely visible. We then manually labeled the superior ventricles on the respective blockface images. All histology ventricle labels were registered to the blockface using the pre-computed registration maps. We computed the 2d DC for all registered histology/block label pairs. The same procedure was performed for our two cases. Mean DC values for case #1 were 0.9082±0.053 (AT8), 0.9112±0.0405 (AT100) and 0.88±0.0531 (MC1). Mean DC values for case #2 were 0.8185±0.1070 (AT8), 0.8385±0.1444 (AT100) and 0.8605±0.1823 (MC1).

#### MRI to blockface 3d registration evaluation

here, we used the superior ventricle masks created by the Freesurfer pipeline (see Methods) and the ground truth MRI labels and were, in turn, registered to the blockface volume using the pre-computed 3d registration files. The DC values were then computed between each of the blockface ventricle label created for the 2d registration evaluation and its respective MRI ventricle label (after registration). The same procedure was performed for our two cases. We obtained a mean DC of 0.9047 (±0.032) for case #1 and mean DC of 0.8570 (±0.095) for case #2.

## Discussion

Molecular imaging is the most promising technique for spatially quantifying proteins in the living brain and has the potential to leverage reliable pre-clinical diagnosis and monitoring progression of neurodegenerative diseases and other brain conditions. A growing number of neuroimaging tracers for ND were developed in the past years, but their bidding properties are not completely mapped in living human brains. Validation of such tracers has been performed mostly using autoradiography assays on tissue sampled from a few brain regions. This approach makes it difficult to understand the nature of off-target signal as well as tracer sensitivity and specificity across different brain areas in living conditions. PET validation using histology data of postmortem brains proved valuable both for beta-amyloid and tau tracers. However, such studies were limited to a few brain areas, and voxel-to-voxel correspondence was compromised by difficulties in registering deformed histological images back to the MRI/PET space. Tracer validation using histopathology data is valuable to PETβ. Creation of whole-brain histological maps for PET tracer validation will leverage the validation of existing tracers by unveiling the biding properties and nature of off-target signal and facilitate the development of new ones.

We developed a scalable, parallelizable pipeline for mapping particles of interest (here, abnormal tau protein) labeled with immunohistochemistry, in whole human brain histological slides by incorporating deep learning algorithms and HPC capabilities in previously developed framework to register histology to neuroimaging. We used this pipeline to create 3D tau density maps that were registered to their respective MRI volume to enable voxel to voxel comparisons. Such a framework can be adapted to any neuroimaging modality such as PET. Previously, we generated histological and computational algorithm to process human brains in the whole that minimizes deformations and facilitate high precision 3D histology to neuroimaging registration. For completing the pipeline, we had to overcome main significant challenges: (1) how to label proteins of interest in supersized (6 × 4”) histological sections, 2) how to image supersized histological slides at high resolution that do not fit regular microscopes; (3) how to perform large-scale image segmentation on several hundreds of images comprising terabytes of data with good accuracy.

We addressed the second challenge by engineering a cost-effective whole slide imaging (WSI) system with a large travel range capable of imaging the entire 4” x6” area of our slides. There has been an explosion in the development of open-hardware microscopy equipment for a wide range of applications [27–31]; with projects relying on a wide variety of materials, from inexpensive 3d printed parts to high-end optical kits. However, those designed focused on small samples. We optimized our scanner to deal with very large tissue samples. As ideal hardware would be economically impractical, we selected the configuration based on the cost-benefit factor for this project. We chose a machine vision objective because it incorporates all the optical elements necessary for microscopic imaging in one lens set that could be attached to the camera, meanwhile providing a resonable view of view side. It optimized scanning time. As it is, it took around 50 to 90 min for our scanner to cover one whole section (depending on the section size), meaning it takes about 233 h to scan a set of sections stained at 1 mm interval, not counting loading slides in and out, focusing and transferring data. Any adjustment (resolution, extra z layers) would increase acquisition time. The availability of open-source software was imperative for the success of our design.

The third challenge was the most complex to solve. Thresholding algorithms to discriminate tau from background failed (see results). Next, we tested available machine learning algorithms, such as Weka, that also failed. Signal to noise ratio in histological images, mainly when obtained from large and/or thick sections tend to be low. Post-processing such images may inset biases or mask real signal. Therefore, we took the challenge to develop a CNN to efficiently segment signal from a background in raw histological images. We chose to use a DL algorithm due to its ability to automatically discover unknown patterns that best characterize a raw data set. DL models data in a bottom-up approach, characterizing it by small low-level features such as edges, in lower network layers and increases in abstraction and complexity in the following layers, thus allowing the model to better capture semantic meaning [32], as opposed to traditional low-level segmentation methods, like thresholding, that is only capable of working at the pixel level. This makes DL more robust to IHC artifacts, such as staining inhomogeneity – where stain intensity changes from slice to slice, and present of off-target stained structures.

The literature on computational methods for locating ND-related proteinopathies is scarce and unsuitable for large datasets generated by large tissue specimens [33–36]. We manage to work with a large volume of data by harnessing the power of HPC to run our segmentation pipeline in an embarrassingly parallel way, where hundreds of copies of the same pipeline were run at the same time.

To test the robustness of our network, we conducted a thorough validation of our results. On ROC AUC, we obtained classification results (mean 0.88 on the testing set and 0.87 on the validation set) are just below the reported values on systems for cell culture [37] (0.95). Noteworthy cell culture images have a clearer background and fewer artifacts than histological images. When compared to computational methods for astrocyte detection in digital pathology [38], which is a more challenging segmentation problem since digital pathology images usually have cluttered background, our pipeline also yielded mean precision (0.88 for testing and 0.92 for validation) above the reported value but much worse recall performance (0.16 for testing, 0.13 validation) when compared to the reported value of 0.78. We see a similar phenomenon when comparing our results to literature in automatic tauopathy segmentation [34]. Our precision results are well above the 0.72 reported but our recall is way below the value of 0.92 reported. The low recall values could only be improved by increasing the resolution of the scanned images, which would require much more scanning time, storage needs, and computing processing (much larger dataset) as neuropil threads are thin and convoluted, making their imaging blur. Detecting blurry objects is a commonly known limitation of CNN

The biggest challenge we faced while working with whole human brain IHC was batch inhomogeneity, where adjacent slides may have different marker concentrations due to variations during IHC staining. IHC inhomogeneity does not seem to greatly impact studies with small tissue sample size but is detrimental to studies with wide tissue areas, like ours. We tackled the problem using a biologically consistent protein concentration normalization procedure, where regions of high and low heatmap signal were selected from high and low stain quality slides during manual screening. The signal was them boosted in low quality stain slides by adding an *increase factor* computed from the reference slides. Another big challenge we faced during this project development was the use of manual affine registration during the 2d histology to blockface alignment (Figure 1, 2b). Although this solution is robust to histology artifacts, it impairs the scalability of our pipeline.

Here we proposed a computational pipeline that takes advantage of a modern deep learning algorithm to create whole-brain protein maps that can be used to validate neuroimaging modalities, overcoming the limitation of in-vitro assays. We used our pipeline to create quantitative tau protein maps since tau is a well-known AD hallmark and highly correlated with clinical decline. The use of deep learning, however, make our pipeline very flexible and easy to retrain to work with other makers, such as

Our 3D mapping at microscopic resolution coupled with our previously developed 3D registration algorithms for combining histological and imaging volumes can potentially open avenues for thorough and systematic validation of new neuroimaging tracers.

## Materials and Methods

### Specimen Procurement, Histological Processing and brain slabbing

We tested the proposed pipeline using two whole human brains. The first specimen, denominated Case #1, belonged to a 88 years old individual, cognitively normal, diagnosed with Braak stage 4 and an A2,B2, C1 score for AD neuropathologic changes [15, 39] and the second, denominated Case #2, belong to a 76 years old patient diagnosed with dementia and Braak stage 6 and an A3, B3, C3 score for AD neuropathologic changes. Case #1 underwent postmortem MPRAGE MRI acquisition while Case #2 underwent postmortem SPGE acquisition within 10 hours of the time of death. Both cases also underwent CT acquisition. After procurement, specimens were fixed by immersion in 4% buffered paraformaldehyde for three weeks. To avoid deformation, we stored the specimens upside down, hanging by the circle of Willis for three days and after that, mounted them in plastic skulls 3d-printed from patient’s CT images and continued fixation for another 18 days inside a bucket filled with buffered 4% paraformaldehyde. After fixation, the specimens were embedded in celloidin, and the blocks allowed to congeal. The blocks were sectioned using a sliding microtome (AO 880, American Optical, USA) equipped with a 14” long C-knife. The brains were sectioned in serial sets of coronal slides, each set containing five 160 μm-thick sections to obtain an accurately aligned stack of unstained serial section photographs, digital photographs were acquired directly from the blockface following each stroke During block sectioning using a high definition DSLR camera (EOS 5D Mark II, Canon, Tokyo, Japan) mounted on a copy stand arm (Kaiser Fototechnik, Germany) and linked to a computer. PET scan voxel size varies from 3 to 5 mm. During the process, brain tissue shrinks about 30 to 40%. Thus, we chose to have five slides in each set, so each PET scan voxel would contain at least two full histological sets in the Z-axis, opening the opportunity to probe multiple markers per voxel. Histological sections were stored in 70% alcohol for further processing. Details for tissue processing celloidin embedding and cutting have been debrided by us elsewhere [13, 40]theofilas 2014}.

### Immunohistochemical labeling of abnormal tau protein inclusions

In Alzheimer’s disease, tau protein becomes hyperphosphorylated and aggregated into inclusions. It is unclear which of these abnormal tau forms the tracers are binding to. We designed our protocol to accommodate different probing forms of protein accumulations. Immunostaining remains an attractive validating method to probe proteinaceous inclusion while preserving tissue integrity and providing spatial information. For this study, we chose two widely used antibodies against phospho-tau, AT100, and AT8. As a rule, all 1^st^ sections from each set were immunostained to detect tau protein phosphorylated at epitopes pSer202+Thr205 (AT8, Thermo). All 2^nd^ sections from even-numbered sets were immunostained to detect tau protein phosphorylated at epitopes pThr212+Ser214 (AT100, Thermo). Immunohistochemistry reactions were done in batches of 50 to ensure homogeneity. Each batch contained positive and negative control for comparison. The controls were serial sections from a single subject, not belonging to the study for quality control purposes.

For the quality verification process, LTG visually reviewed the positive control slide of each batch and compared the staining process to other tissue stained with the same antibody to assess the adequacy of the staining process. Staining was deemed unacceptable if no staining was present, very weak staining was observed as compared to another stained positive control slides, and/or very strong background, like the level of staining, was observed. Batches whose positive control was deemed unacceptable were excluded, and the staining process was repeated on the 2^nd^ set of contiguous tissue sections, which in turn went through the same quality control process.

### Whole Slide Imaging

We built a cost-effective whole slide scanner (Figure 3) to accommodate our histological slides (5” × 6”), which do not fit into regular microscope stages or cannot be fully imaged due to short stage travel range. The hardware is comprised of a high-precision, 6” travel range, industrial XY stage (Griffin Motion), an Olympus manual focusing box, color CCD camera (Qimaging Micro publisher 6) and 5.5X machine vision objective (Navitar Zoom 6000) mounted directly on the camera. Illumination is performed by a lightbox with diffuser, mounted on top of the XY stage. Slides are loaded directly to a 3d printed slide mount fixed on top of the lightbox. The scanner is controlled by a software developed in-house using Macro Manager 2.0^42^, which has a user interface that allows the user to select the region-of-interest (ROI) to be scanned and white balance and lens magnification parameters. It computes the coordinates of the image tiles necessary to cover the selected ROI and is responsible for synchronizing the XY stage movements with image capture. The full resolution histological images (example in Figure 2) is created by *stitching* the tiles together using TeraStitcher^19^, which is capable of working with several Gigabytes of data while maintaining a small memory footprint. During the stitching process a 10% resolution version of the image is also created and is used during the histology pre-processing and registration steps and also for visual inspection. This setup yielded a 1.22um pixel resolution. We release our scanner software can be downloaded in (https://github.com/mary-alegro/LargeSlideScan).

### Creation of datasets for SlideNet training and validation

In order to create our training, testing, and validation datasets, we drew 1024×1024 pixel patches from randomly selected gray matter locations throughout the entire full-resolution brain image dataset. A set of 100 patches was created for each marker (*AT100* and *AT8*), from each brain, respectively, totaling four datasets. The patch extraction routine was written in Python and completely automated, running on UCSFs’ Wynton cluster in an embarrassingly parallel way. https://wynton.ucsf.edu

Each patch was manually masked for background and tau inclusion with the help of Fiji’s Trainable Weka Segmentation plugin. Here, the user manually selected sample pixels belonging to *tau* and background classes that were, in turn, used to compute several features such as Gaussian filters, Hessian, membrane projections, mean, maximum, anisotropic diffusion, Lipschitz, Gabor, Laplacian, entropy, Sobel, a difference of Gaussians, variance, minimum, median, bilateral filter, Kuwahara, derivatives, structure, and neighbor values. A linear SVM classifier (LibLinear) was then used to generate an initial tau segmentation. The user would retrain and refine the initial segmentation until they obtained satisfactory results. Masks where manually fine-tuned using an image editor (Gimp). All final masks went thought quality control by a highly experienced pathologist.

Finally, patches for each marker (AT8, AT100, and MC1) were combined on bigger datasets, totaling two datasets of 200 patches each and were randomly split in 80% of patches for training, 10% for testing and the remaining for validation.

### Histology Pipeline

#### Pre-processing blockface images

Blockface images have their background segmented using a semi-automated graph-based algorithm. Briefly, the user selects brain and background sample pixels using a graphical user interface (GUI). Images are then converted to LAB color space, and the algorithm computes mean color difference maps (ΔE). ΔE is defined as the distance in LAB space:

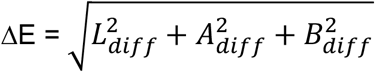

With *L*_*diff*_, *A*_*diff*_,*B*_*diff*_ being the difference values computed as:

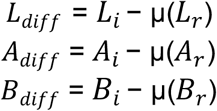

Where *L*_*i*_, *A*_*i*_ and *B*_*i*_ are image LAB channels, *L*_*r*_, *A*_*r*_, *B*_*r*_ are LAB channels of reference pixels selected by the user, and μ(.) is the mean. Pixels in ΔE, whose color is similar to the reference values, appear dark in (smaller distance) while cell pixels are brighter (larger distance). We compute brain and background ΔE maps using the manually selected pixels as reference values and perform a global? Histogram threshold using the Otsu’s method to obtain binary masks. We, in turn, combine both masks to obtain the brain segmentation.

It is common that several undesired objects linger after the initial segmentation. It is usually caused by reflections from the liquid on top of the celloidin block. The segmentation is further refined using a graph-based method to remove the undesired objects. In this method, image objects and their relationship are modeled as a weighted graph, where connected structures are considered the vertices. Edge weights are computed using a similarity function computed from color and distance values. The graph is partitioned using NCuts, leaving just the brain area.

Commonly, the camera or brain must be repositioned several times during sectioning to adjust for changes in block size, causing the blockface images to be misaligned in relation to each other. We use semi-automatic registration to correct this artifact. Having the middle slice as the reference image, we use Matlab’s’ registration GUI to select landmarks for computing affine registrations. Finally, the aligned blockface images are stacked together to form the blockface 3d volume using ITK. This 3d volume is later used as an intermediate space for creating the 3d tau-protein maps.

#### Low-res histology background segmentation

The 10% resolution histological image is converted to LAB color space. In that space, background pixels are consistently darker than brain pixels, and segmentation is performed through histogram thresholding using the triangle algorithm. The resulting binary masks, here denominated brain masks, are used to erase all background pixels. Moreover, brain masks are combined with white matter mask to create gray matter masks that guide the entire segmentation process and are also used during patch extraction for generating our training and validation datasets.

#### 2d registration to blockface image

After background segmentation, the 10% resolution histological images are aligned to their respective blockface images using a combination of manual and automatic registration. Due to the presence of an excessive number of artifacts caused by histological processing, such as tearing, shrinking and shearing, we decided for a manual initialization of the registration using MIPAV spline-based registration. Here, the user manually selects landmarks on both the histology and blockface images. MIPAV then generates a warped image and a registration warp file. After the initial registration, the image goes through a diffeomorphic registration using the 2d SyN algorithm, which is based on the large diffeomorphic deformation model metric mapping (LDDMM) method. Both the registered images and registration mappings are saved for use later in the pipeline.

#### Image and Mask Tiling

Here, each full-resolution histological image is first tiled to reduce memory footprint during image segmentation, with tile size corresponding to approximately to 5mm^2^ of tissue. Tile coordinates and dimensions are saved as XML metadata files. The histological images’ respective 10% resolution gray matter masks are then rescaled to match their full-resolution dimension and tiles following the same procedure used for histology. Finally, histological tiles are masked using their respective gray matter tiles, leaving only the regions of interest on the image. Tiles that are left empty or having less than 5% of tissue pixels are ignored during segmentation. This way, we reduce the overall computational time and guarantee we are only processing relevant information. Image and mask tiling routines were developed in Python and ran on UCSFs’ Wynton cluster in an embarrassingly parallel way, having one pipeline instance for each histological image.

### Deep learning-based segmentation

A convolutional neural network is the core of our segmentation pipeline. We designed SlideNet, a UNet based neural network capable of working with color information and outputting tau presence confidence maps that are later thresholded to create binary maps.

#### Model development and training

Our model has an 204×204×3 pixels input layer to accommodate RGB images – we chose this size to match 1mm^2^ of tissue in our WSI scanner resolution. In our model, the image is pushed through three contractions blocks, the bottleneck, and upsampled by three expansion blocks. Figure 3 shows the SlideNet architecture together with each layer tensor size.

Each contraction block is composed of two convolution layers that use 3×3 kernels, stride of 1 and ReLu activation, followed by a 2×2 max pooling layer and 0.1 rate dropout. The bottleneck is composed of two convolutional layers that use 1×1 kernels, stride of 1 and ReLu activation. Expansion blocks are composed of a 2×2 upsampling layer followed by a 0.1 rate dropout and two convolutional layers that use 3×3 kernels, stride of 1 and ReLu, except for the last expansion block that use 3×3 upsampling. The last layer reshapes the data to a 20000×2 tensor and softmax activation.

Training is performed using standard backpropagation, with cross entropy as the loss function and Adam for optimization. The learning rate is estimated using the cyclical learning rate method [24]. The network was developed in Keras (https://keras.io) on top of TensorFlow (https://www.tensorflow.org). We trained the network using minibatches of 32 images during 100 epochs, or until the loss curve plateaued. We also used massive data augmentation, performing real-time random rotations, shear, horizontal and vertical flips. All training and inference were performed on a NVIDIA Titan V GPU with 12GB of RAM.

SlideNet outputs confidence maps of the existence of tau, which are thresholded to generate binary masks. The threshold value was set to 0.7 for both AT100 and AT8 datasets and pixel with confidence higher than this value was considered tau.

### Heatmap computation

The binary tiles are transferred back to our cluster, where the heatmaps are computed. On each tile, we compute the mean amount of tau, indicated by the mean number of pixels belonging to foreground, inside a 1 μm^2^ of tissue, roughly an 8×8 pixels block. The heatmaps are generated as tiles having the same dimension of binary tiles where each 1 μm^2^ block is filed out with the mean number of pixels belonging to tau. Tiles are stitched together to create a heatmap having the same resolution as the original histological image and is latter resized to 10% of its resolution.

#### Heatmap normalization

Each of the four heatmap datasets (AT100 and AT8 for each brain) are normalized to mitigate batch staining inhomogeneity. For each dataset, a trained neuropathologist selects strong signal ROIs from slices whose staining is deemed optimal, and weak signal ROIs from slices whose staining is suboptimal. Outliers are removed from each ROI by thresholding the values are the first and third quartiles and mean strong and weak signals are calculated. We then compute an *increase factor* as the absolute difference value between both means. All heatmaps from slices that were deemed suboptimal during quality control are adjusted by summing their non-zero values with this *increase factor*.

#### Heatmap to blockface alignment and stacking

The 2d registration maps computed during the 2d registration to blockface image step are applied to the 10% resolution normalized heatmaps, yielding heatmap registered to their respective blockface images. These images are then stacked together to for a 3d volume.

### MRI acquisition and processing

Postmortem structural T1-weighted MR imaging of Case 1 was performed on a 3T Siemens Skyra MRI system with a transmit and 32-channel receive coil using a 3D MPRAGE T1-weighted sequence with the following parameters: TR/TE/TI = 2300/2.98/900ms, 176 sagittal slices, within plane FOV = 256×240mm^2^, voxel size = 1×1×1mm^3^, flip angle = 9°, bandwidth = 240Hz/pix.

Postmortem structural T1-weighted MR imaging of Case 2 was performed on a GE Discovery 3T MR750 system with a transmit and 32-channel receive coil using a 3D SPGR T1-weighted sequence with the following parameters: TI = 400ms, 200 sagittal slices, within plane FOV = 256×256mm^2^, voxel size = 1×1×1mm^3^, flip angle = 11°, bandwidth = 31.25Hz/pix.

Accuracy of medical image registration approaches relies presence and detection of homologous features in both target image (i.e., 3D blockface reconstruction) and the spatially warped image (i.e., structural T1-weighted MRI). To satisfy this prerequisite, using the Advanced Normalization Tools (ANTs) [41], the N4 non-parametric non-uniform intensity normalization bias correction function [42, 43] followed by skull-stripping was applied on the structural T1-weighted MRIs. Briefly, structural T1-weighted MRIs were warped into the 3D image space of the corresponding blockface reconstructions using ANTs [21, 22]). First, a linear rigid transformation was performed. Then a diffeomorphic transformation using the Symmetric Normalization (SyN) transformation model was performed. SyN uses a gradient-based iterative convergence using diffeomorphisms to converge on an optimal solution based on a similarity metric (e.g., cross-correlation) [44]. To validate the registration we calculated the dice of ventricles.

### Histology to MRI 3D registration

Three-dimensional histology to MRI registration is performed using the 3D SyN algorithm. SyN is based on the large deformation diffeomorphic deformation model metric mapping (LDDMM) method whose mathematical properties are especially appealing for our problem. Diffeomorphic maps are smooth and invertible functions guaranteeing that no folds or vanishing tissue will occur during the registration process. These models also enforce a 1-to-1 mapping between the movable and reference voxels and are suitable to handle large deformations. SyN is a mature method that has been reported to outperform several popular registration methods. Here, SyN is used with Mattes mutual information for measuring similarity, given that it is the most robust option for handling multi-modality registration problems. All 3D registration maps computed in this step are stored for future use. Figure 3h is an example of 3D histology to MRI registration using the described method. The first row shows sagittal, axial, and coronal checkerboard representations of a histology slice, overlaid on its respective MRI. The right-hand side shows a 3D rendering of one registered volume overlaid on its MRI.

### 3d Reconstruction and Visualization

The 3d registration maps generated during the <u>*Histology to MRI 3D registration*</u> are inverted and used to warp the MRI to blockface space, allowing direct comparison between MRI signal and tau density. Freeview is used for visualization of 2d slices. Amira (ThermoFisher Scientific) is used for 3d reconstruction and visualization.

